# Modelling the effectiveness of targeting Rift Valley fever virus vaccination using imperfect network information

**DOI:** 10.1101/2022.10.04.510793

**Authors:** Tijani A. Sulaimon, Gemma L. Chaters, Obed M. Nyasebwa, Emanuel S. Swai, Sarah Cleaveland, Jessica Enright, Rowland R. Kao, Paul C. D. Johnson

**Affiliations:** The Roslin Institute, University of Edinburgh, Easter Bush Campus, Midlothian, Scotland, United Kingdom; School of Biodiversity, One Health and Veterinary Medicine, University of Glasgow, Glasgow, Scotland, United Kingdom; School of Computing Science, University of Glasgow, Glasgow, Scotland, United Kingdom; Royal (Dick) School of Veterinary Studies, University of Edinburgh, Easter Bush Campus, Midlothian, Scotland, United Kingdom; Institute of Infection, Veterinary and Ecological Sciences, University of Liverpool, Liverpool, United Kingdom; Veterinary Council of Tanzania, Dodoma, United Republic of Tanzania; Department of Veterinary Services, Dodoma, United Republic of Tanzania; Anicare Vet Services, Tanga, United Republic of Tanzania

**Keywords:** Tanzania, livestock networks, network measures, metapopulation model, Rift Valley fever, targeted vaccination, imperfect information, robustness

## Abstract

Livestock movements contribute to the spread of several infectious diseases. Data on livestock movements can therefore be harnessed to guide policy on targeted interventions for controlling infectious livestock diseases, including Rift Valley fever (RVF) — a vaccine-preventable arboviral fever. While detailed livestock movement data are available in many countries, such data are generally lacking in others, including many in East Africa, where multiple RVF outbreaks have been reported in recent years. Available movement data are imperfect, and the impact of imperfect movement data on targeted vaccination is not fully understood. Here, we used a network simulation model to describe the spread of RVF within and between 398 wards in northern Tanzania connected by cattle movements, on which we evaluated the impact of targeting vaccination using imperfect movement data. We show that pre-emptive vaccination guided by only market movement permit data could prevent large outbreaks. Targeted control (either by the risk of RVF introduction or onward transmission) at any level of imperfect movement information is preferred over random vaccination, and any improvement in information reliability is advantageous to their effectiveness. Our modelling approach demonstrates how targeted interventions can be carefully applied to inform animal and public health policies on disease control planning in settings where detailed data on livestock movements are unavailable or imperfect due to a lack of data-gathering resources.

## 1 INTRODUCTION

Infectious diseases are an important challenge facing livestock production systems (1) particularly in developing countries, due to their substantial impact on livestock health and welfare, and in terms of economic losses (2, 3). Worldwide, infectious diseases of animals that affect humans (zoonoses) are responsible for over 2.5 billion cases of illnesses in humans, with an estimated 2.7 million deaths every year (4, 5). Due to the impacts of many factors — including climate change, urbanisation, globalisation, changing eating habits, deforestation, and human-wildlife interaction — human and animal populations are at high and increasing risk of zoonotic disease transmission, emergence, and re-emergence (6, 7, 8).

Infectious diseases, including zoonoses, can spread within and between livestock populations by various routes, including direct contact, via vectors such as mosquitoes, consumption of contaminated animal products, and livestock movements (9). Livestock movements, for trading and grazing, play a significant role in disease spread between populations (10). For instance, the 2001 foot-and-mouth (FMD) outbreak in the UK was primarily driven by long-distance movements of livestock between holdings and local transmission within holdings at the early stage of the epidemic before a movement ban was implemented (11, 12).

Vaccination is an effective measure for controlling infectious disease outbreaks (4). However, to design an efficient vaccination strategy for infectious disease control, it is crucial to understand the behaviour of disease transmission within and between populations. Decisions about where to impose a disease control strategy rely on a range of factors, including the specific pathogen, the outbreak type (whether it is a common source or propagated outbreak), the size of the target population, the contact network connecting the population, the resources available, and the effectiveness of the control strategy. Network-based modelling is a way of describing the interplay between infectious disease transmission and the contact network pattern, as well as providing a means of testing the impact of interventions in silico (13, 14). Targeted vaccination strategies that exploit the hierarchy of nodes’ connectivity in a network are highly effective when the complete structure of the network is known (14, 15, 16, 17). However, our knowledge of the network structure of populations is usually imperfect due to incomplete or unreliable data (18, 19). In reality, network studies rely on observed networks that differ from the true network connecting the population under investigation (19). Such error can substantially impact network measures, which might considerably affect their performance in epidemic control on the true network (19, 20), therefore motivating analyses to understand how different network measures are impacted by imperfect information.

Tanzania is one of the countries in East Africa where infectious diseases, including zoonoses, pose a challenge to livestock and human health and welfare. Several efforts have been made to reduce the burden of dangerous zoonotic diseases, including rabies, Rift Valley fever (RVF), and brucellosis in Tanzania (21). For example, in 2010, the Tanzanian government launched a nationwide vaccination campaign against rabies, following many other countries aiming to meet the global target for eliminating human deaths caused by rabies by 2030 (22). As resources are often limited in low-income countries such as Tanzania, insights from modelling analyses can help guide such vaccination campaigns to yield the greatest impact on disease outbreaks.

Livestock are typically managed in distinct units, which in Tanzania comprise herds and flocks that are kept in households or multi-family compounds within villages. Livestock units are linked by movements, many of which go through markets (23, 24), forming a complex network of nodes (representing populations of livestock) and links (representing livestock movements between populations). Such networks can be modelled using a meta-population framework, which allows us to test the impact of targeted vaccination strategies (13). Many researchers have investigated the impact of network-based vaccination strategies on epidemic spread (14, 13). Many of those studies are either based on theoretical (model) networks (16, 15, 17, 25, 26, 27) or real-world networks generated from developed countries, where livestock movement data are rich (28, 10, 29, 30, 31). So far, however, few studies have used network structure in African livestock setting to study vaccination strategies for infectious disease control (24, 32, 33, 34, 35).

Here, we investigate the effectiveness and robustness of various vaccination strategies under imperfect knowledge of network structure. To do this, we simulated the spread of RVF virus using a stochastic Susceptible-Exposed-Infected-Recovered (SEIR) model on both theoretical networks (generated by simple network models, including random, scale-free and small-world) and a network of livestock movements across three regions in northern Tanzania (24), upon which we tested the impact of simulated targeted vaccination. Theoretical networks were included to test the robustness of our conclusions to network structure. We focused on RVF as an exemplar because it is an important vaccine-preventable livestock infection where we can build on previous studies of the role of livestock movements in the spread of infection (24). The aims of this study were: first, to model the effectiveness of epidemic control using network-informed vaccination strategies; and second, to investigate how their performance varies under imperfect knowledge of the network.

## 2 METHODS AND MATERIALS

### 2.1 Overview

This *in silico* study followed five basic steps: 1) generate a cattle movement network; 2) simulate the condition of imperfect information about the network; 3) use different network-targeted strategies to select nodes for vaccination under imperfect network information; 4) simulate multiple scenarios of Rift Valley fever (RVF) virus transmission over the network with selected vaccinated nodes; and 5) compare mean outbreak size after one year under various vaccination strategies and levels of imperfect network information. We begin this section by describing the study area and how cattle movement networks were generated, followed by a description of the network-based vaccination targeting strategies (using degree, betweenness, and PageRank centrality measures) and risk-based strategies (the risk of RVF introduction into the cattle population, derived from an index for indicating rainfall) that were considered in this study. The last three subsections focus on the RVF virus transmission model, how vaccination was implemented, and how scenarios of imperfect network information were modelled.

### 2.2 Study area

The study area comprises three regions — Arusha, Manyara, and Kilimanjaro — located in the northern part of Tanzania, containing approximately 4.8 million inhabitants in a total area of 95 348 km^2^ (36). Livestock production in these regions is predominantly carried out for food, income, and social status (24, 23). A detailed description of the study area and its livestock practices is given in de Glanville et al. (23). There are various complex socio-economic reasons for livestock movement in northern Tanzania, including access to natural resources such as grazing, water, and salt (37), exchange of livestock as gifts and payments (38), and trade movements through the market system (24, 39).

### 2.3 Network generation

This study exploits the multiplex network of cattle movements in northern Tanzania generated by Chaters et al. (24). The multiplex (which we shall refer to here as the “data-driven” network, see Figure 2) contains two layers: a movement network of cattle through markets and a network connecting adjacent wards. In the first layer, Chaters et al. (24) market movement permit data were used to generate a static network of cattle movements between 398 wards within three regions (Arusha, Kilimanjaro, and Manyara) in northern Tanzania, where the number of cattle moved is based on the estimates of the number moved in a month. A *ward* is an administrative unit of a mean area 243 km^2^ containing a mean human population of 12 000 and a mean cattle population of 9 000 across all 398 wards (24). The market permit data were collected as part of the SEEDZ (Social, Economic and Environmental Drivers of Zoonoses in Tanzania) project, the protocols and procedures of which were approved by the ethics review committees of the Kilimanjaro Christian Medical Centre (KCMC/832) and National Institute of Medical Research (NIMR/2028) in Tanzania, and in the UK by the ethics review committee of the College of Medical, Veterinary and Life Sciences at the University of Glasgow (39a/15). Approval for study activities was also provided by the Tanzanian Commission for Science and Technology (COSTECH) and by the Tanzanian Ministry of Livestock and Fisheries, as well as by regional, district, ward and village-level authorities in the study area.

**Figure 1.**
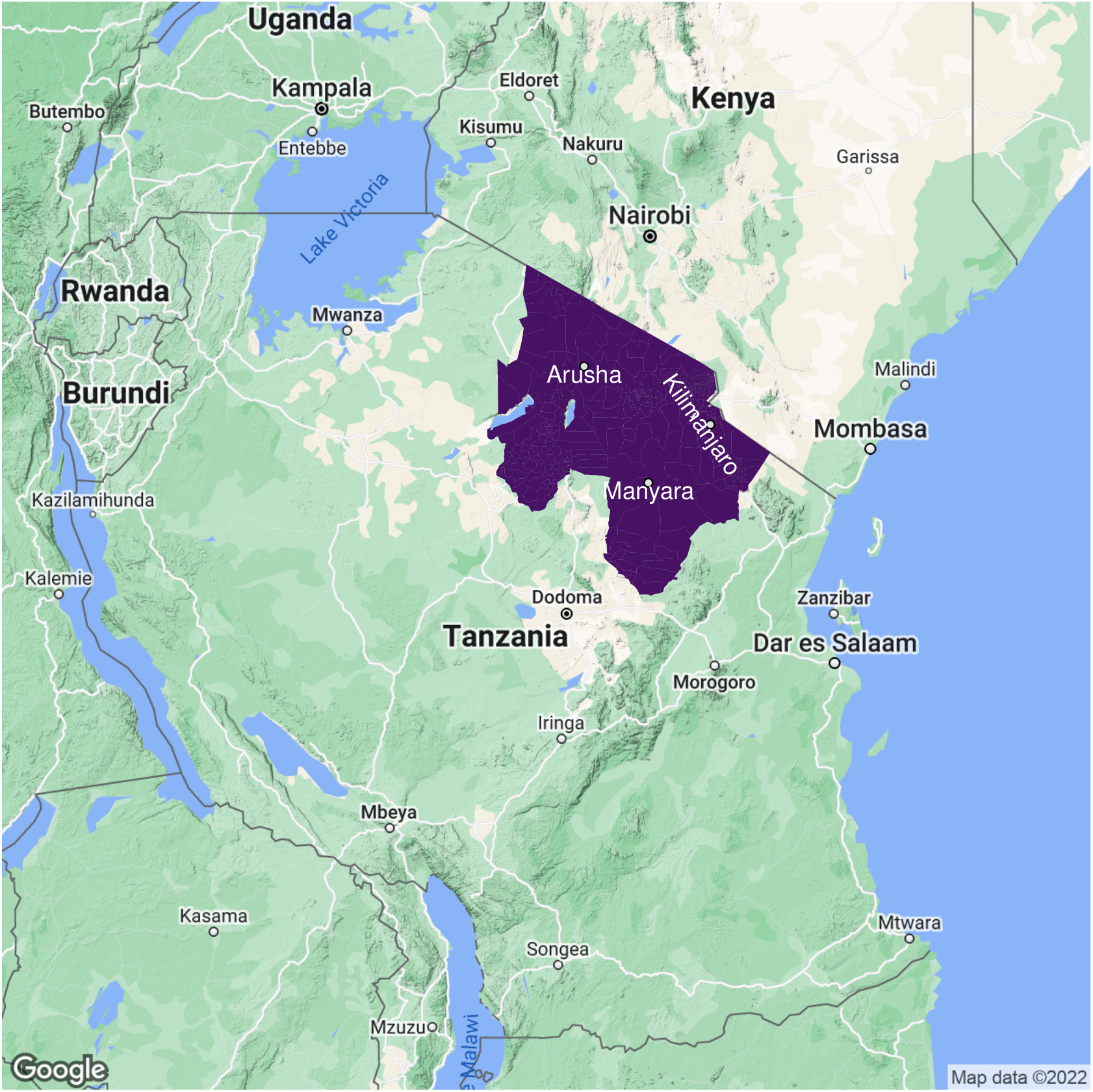
Map of the study area in northern Tanzania. Arusha, Manyara, and Kilimanjaro regions are highlighted in purple. Map created using Google map data available through ggmap package (40) in R. Shape files obtained from NBS, United Republic of Tanzania https://www.nbs.go.tz/index.php/en/census-surveys/gis.

**Figure 2.**
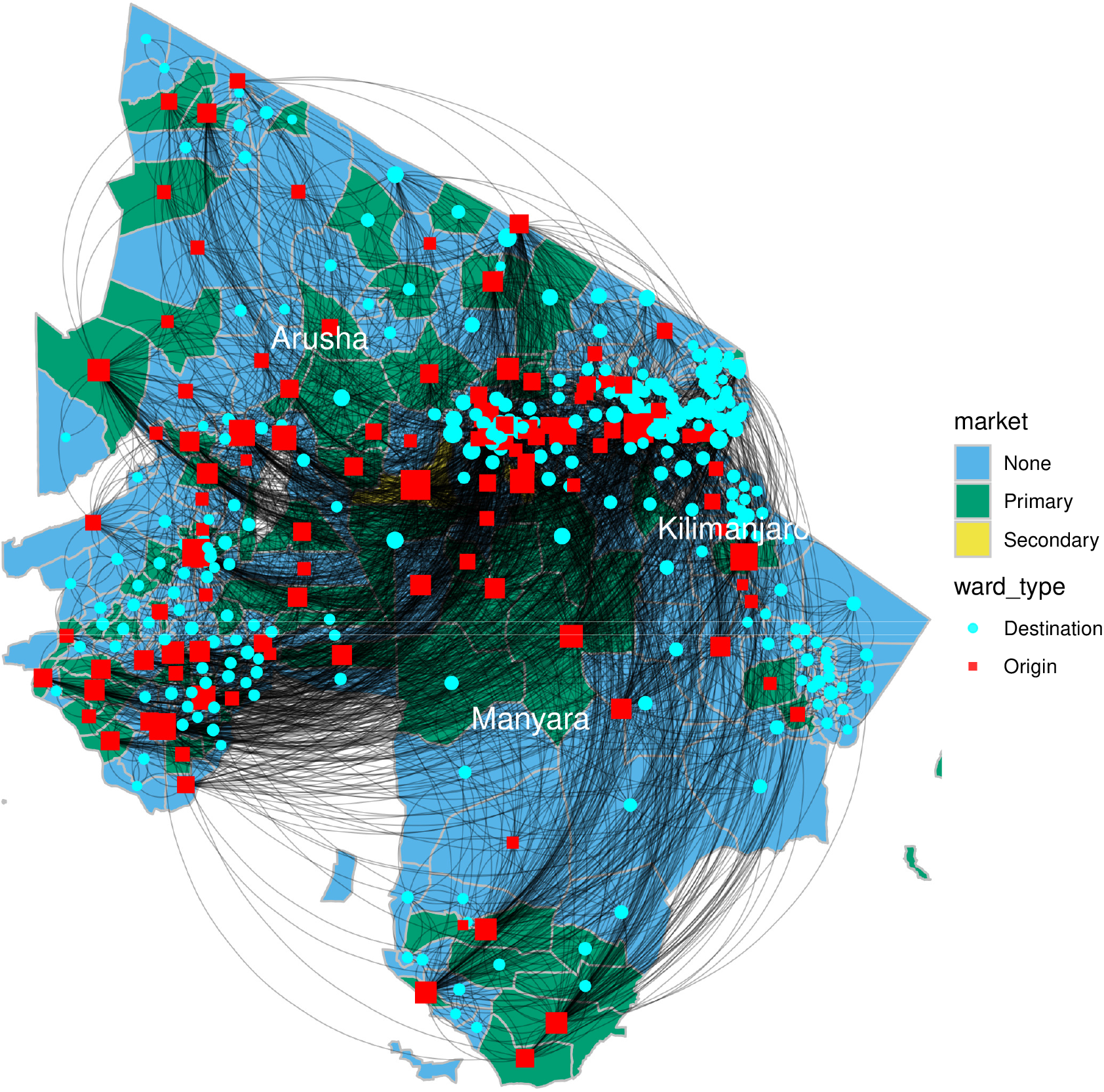
Weighted multiplex network of livestock movements between 398 wards (nodes) in northern Tanzania. Node area is proportional to node degree. Generally, cattle purchased in primary markets are batched into larger groups by traders and transported to secondary markets, where prices are higher. A node is either an origin, a destination, or both. An origin is a node with at least one outward movement of cattle via market. Movements of cattle are predominantly through primary or secondary markets, with secondary markets having the largest volume of trade (weighted degree of nodes). In addition, each ward is connected to adjacent wards through local movements of cattle, representing non-market movements such as private sales and gifting, which were not captured by the movement permit data.

The cattle movement network data were aggregated spatially at the ward level because the destinations recorded on the movement permits could not typically be located at a finer scale (24). A spatial network, created by connecting each ward to all its spatially adjacent wards, was added to the market movement network, as a means of accounting for contacts that occur between wards through sharing of grazing and water sources, and movements of animals through gifting and private sales (24). We used the combined monthly market movement networks to generate a static directed weighted network over a year, which was then used to calculate network measures. The spatial network was excluded from the calculation of network measures because we expect targeting strategies would be guided by market network data in real policy situations.

We generated unweighted theoretical networks with an equal number of nodes (398) and mean degree (4) as the monthly “unweighted mean market movement” network to explore the effect of varying network structures on results. These included Erdős-Rényi (ER) random networks, scale-free networks, and small-world networks. Only the information on whether there was a market movement of cattle between wards in the data-driven network was used to generate the unweighted mean market network.

For random networks, we use the Erdős and Rényi *G*(*N, M*) model with *N* = 398 nodes and *M* = 2×398 links, resulting in random networks with mean degree 4 (41).

To generate small-world networks, we followed the model proposed by Watts and Strogatz. First, arrange nodes (wards) in a 1-dimensional circular lattice (a ring) such that each node is connected to two nearest neighbours on either side. With probability *p* = 0.1 (which is above the percolation threshold), each link is rewired, resulting in long-distance links (shortcuts) (42).

For scale-free networks, we adapted the algorithm described by Albert and Barabási (26). We begin with a fully-connected network generated using the preferential attachment model, with *N* = 398 nodes, *m* = 2 (the number of links added at each time step), and power = 1. The giant strongly connected component (GSCC) size of networks created by this algorithm is generally 1. To obtain networks with a larger GSCC size, we reshuffled the network’s links while preserving the degree distribution. The summary statistics of networks generated are presented in Table 1. All networks are directed and were generated using the ‘igraph’package (43), available within R version 4.1.1 (44).

**Table 1.** Summary statistics of the networks generated. N, number of nodes; ⟨*k*⟩, mean degree; std (deg), standard deviation of network degree; CC, mean clustering coefficient; GWCC and GSCC, giant weakly and strongly connected components. ^†^The data-driven network is weighted. Its unweighted version has a mean degree of 4, like the theoretical networks, which are unweighted.

| Network | N | $\langle k \rangle$ | std(deg) | CC | GWCC | GSCC |
| --- | --- | --- | --- | --- | --- | --- |
| Erdős-Rényi | 398 | 4 | 1.99 | 0.01 | 390 | 253 |
| Scale-free | 398 | 4 | 7.22 | 0.02 | 398 | 295 |
| Small-world | 398 | 4 | 0.87 | 0.25 | 398 | 389 |
| Data-driven<sup>†</sup> | 398 | 14.5 | 11 | 0.30 | 398 | 398 |

### 2.4 Network measures

In a spreading process, the importance of a node in a network is characterised by its structural position, and its contribution to epidemic spread over the network (45). Highly central or influential nodes are likely to infect or expose many other nodes disproportionately, and potentially drive the speed and severity of epidemic outbreaks (46, 47). Identifying the most central nodes is essential for breaking the transmission chain and slowing down the speed at which an epidemic is spreading (16, 17, 15). Here, we study three standard centrality measures (degree, betweenness, and PageRank) widely considered to be relevant to disease spread (48, 49, 50, 16, 51, 15).

The degree centrality of a node in an undirected network is the number of links connected to the node; for directed networks, this may be out-degree or in-degree (52). The degree centrality is vital in studying infectious disease transmission because it measures the number of potentially infectious contacts (13, 52, 17). Betweenness centrality measures the extent to which a node lies on the shortest path connecting other nodes of the network (53). In the context of disease spread, it describes the importance of a node to disease propagation across the communities of the network (13). PageRank is a metric used to rank web pages in the Google search engine (54). It ranks web pages based on the number of backlinks (number of links pointing to the page or in-degrees) or highly important backlinks (54). It indicates the probability of visiting a node by a random walker in a network (54). Each of these measures assigns “importance” differently and has their strengths and shortcomings.

### 2.5 Risk of RVF introduction into cattle population

Mosquitoes are both reservoirs and vectors for the RVF virus and, therefore, capable of keeping the virus in the enzootic cycle for a very long time — even in the absence of livestock — through vertical transmission from infected adult female mosquitoes to their offspring (55). Mosquito-borne diseases such as RVF are susceptible to climate-mediated changes (56, 57). Abnormally high rainfall can trigger hatching and amplification of mosquito vectors, hence provoking outbreaks (58). Several studies have demonstrated that the remotely sensed Normalised Difference Vegetation Index (NDVI), one of the most used indices for green vegetation, is a good indicator of rainfall and conditions suitable for the emergence of RVF (59, 57, 60). According to Linthicum et al. (61), it is possible to anticipate RVF outbreaks in East Africa up to 5 months before they occur using a range of climate indices, including NDVI.

To calculate RVF introduction risk, the average monthly NDVI raster for the study area between January 2000 and December 2017 was divided by wards into grid cells. The risk score of a ward in the data-driven network is the proportion of cells of the ward that has an NDVI value between 0.15 and 0.4 (an indicator of regions with high rainfall, roughly equivalent to mean annual rainfall between 100 and 800 mm) (60). In theoretical networks, risk scores were randomly distributed.

### 2.6 Disease simulation

An SEIR model was used to describe the transmission dynamics of RVF virus within wards. At any time, each bovid belongs to one of the four states: susceptible (S), exposed (E), infectious (I), or recovered (R). Susceptible individuals get infected and moved to the exposed class at rate 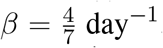. Exposed individuals become infectious at a rate 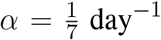 and recover at a rate 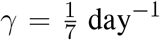. We modelled transmission via vectors as delay in infectiousness onset through the state E as a large-spatial-scale approximation for the vector incubation stage and the latent period in cattle (62). The basic reproduction number, 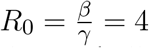, is defined as the expected number of secondary infections produced by an infected individual in a wholly susceptible population (63).

Because it would be unrealistic to assume homogeneous mixing of cattle within a ward, each ward was divided into 64 grid cells in an 8 × 8 matrix of sub-nodes, with each cell representing a sub-node, i.e. a sub-village within each ward. Within each cell, transmission is driven by a homogeneous mixing SEIR model. Transmission between proximal sub-nodes within each ward was allowed through a spatial coupling process to account for the risk of infection transmission between sub-nodes through interactions such as using shared resources. The proportion of infections produced in a sub-node *i* through the coupling process at any time *t* is defined by

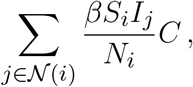

where *N*_*i*_ is the total number of cattle in ward *i, C* = 0.02 is the coupling strength, and 𝒩 (*i*) is the set of the first neighbours of the sub-node *i*.

Cattle were moved between network wards through network links every month (30 days) to allow RVF virus to spread between wards. The network link weights determined the number of cattle moved in the data-driven network. In contrast, cattle movements in the theoretical networks were based on the mean number of animals moved through market. In each simulation scenario, we select a proportion of nodes for vaccination based on the strategy of interest. Since we are only interested in successful epidemics, we seeded infection in five wards based on disease introduction risk (see Figure 3) and with 10 infected cattle in each of those wards, to reduce the probability of stochastic extinction, as less intensive seeding resulted in many failed simulations. For each ward selected for disease introduction, seeded cases were uniformly distributed between grids of the ward, and the epidemic was allowed to spread locally and through cattle movements within and between wards, respectively. The numbers of animals in each state were updated at daily time steps. All simulations were implemented using the SimInf package (64) within the R statistical software environment (44).

**Figure 3.**
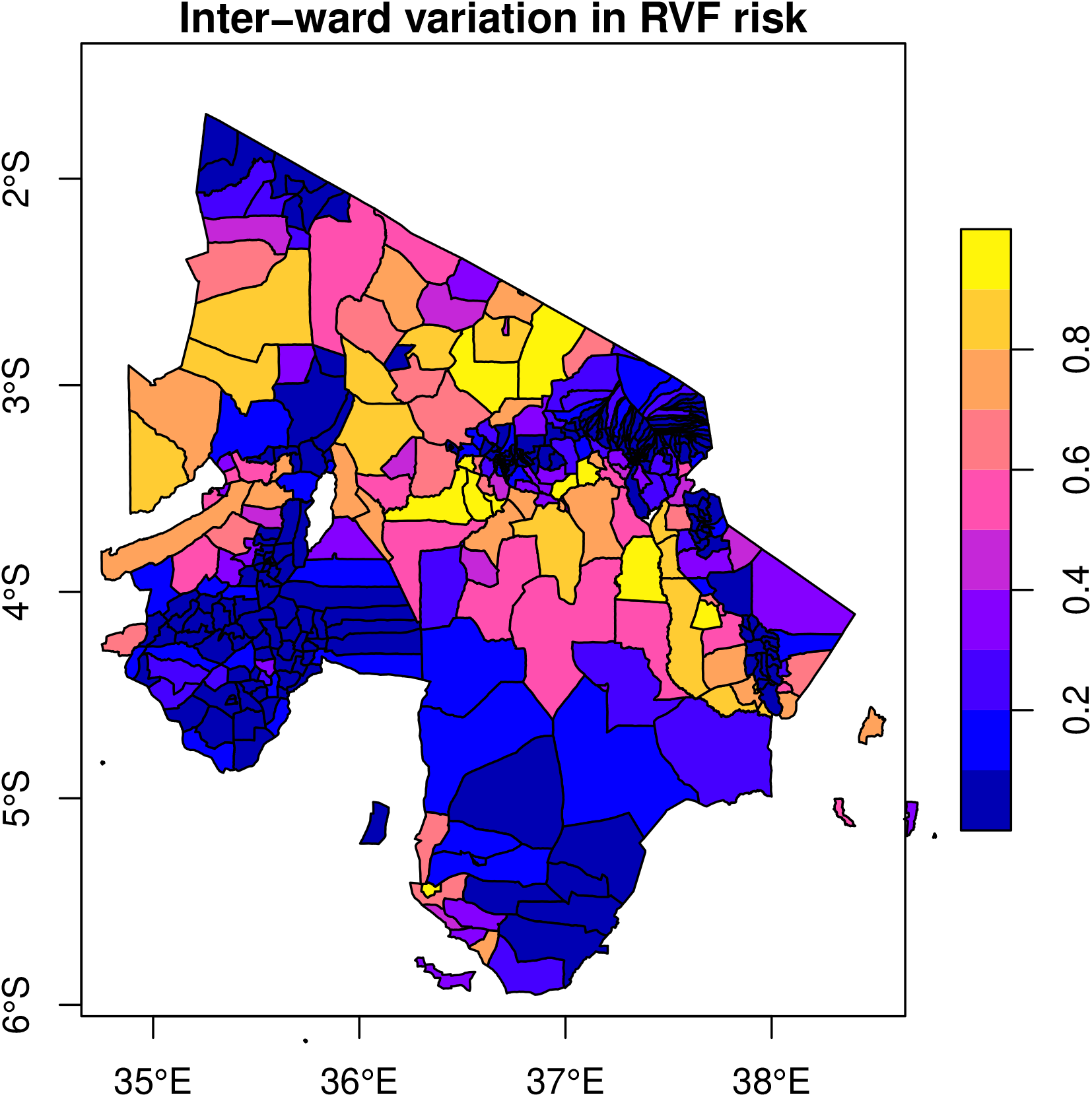
The distribution of RVF emergence risk was simulated by linking disease emergence to climate suitability for RVF vectors. RVF risk score ranges between 0 and 1 (right); a high-risk score represents high RVF risk.

### 2.7 Vaccination strategies

To explore the impact of vaccination on RVF virus transmission dynamics, a proportion of nodes (10%, 20%, 30%, 40%, and 50%) were selected for pre-emptive vaccination (i.e. in anticipation of a higher risk of disease outbreaks), at 75% within-node coverage (the proportion of cattle to be vaccinated in each ward to achieve herd immunity, 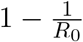). Once vaccinated, each bovid might become immune to the infection with some probability, *e*, which represents the vaccine efficacy. In this model, we assume *e* = 100% (that is, the vaccine is 100% efficacious) and no waning immunity. Immune cattle cannot become infected and do not contribute to disease transmission.

Given limited resources, we wish to select a proportion of nodes for vaccination to ensure the greatest reduction in epidemic size. Here, we considered four vaccination strategies. These included three network-based strategies, where nodes with high degree, betweenness, or PageRank centrality rank are targeted for vaccination, and a risk-based strategy motivated by habitat suitability for disease occurrence (Figure 3). The aim of comparing network-based against risk-based strategies was to understand whether the risk of introduction of disease is a better determinant of outbreak size than the risks associated with propagation across the network. We also considered random vaccination and a no-vaccination scenario to establish two baseline measures for comparison to targeted vaccination scenarios.

The effectiveness of vaccination strategies under perfect information was measured in terms of percentage reduction in the mean number of wards with at least one infected bovid (MNWIC) under each vaccination scenario relative to the MNWIC under the no-vaccination scenario across all 400 simulations. Effectiveness was calculated as (no-vaccination MNWIC minus vaccination MNWIC) / (no-vaccination MNWIC).

### 2.8 Simulating imperfect network information

To examine the effectiveness of vaccination strategies under conditions of imperfect but unbiased cattle movement network data, we used the following process: (1) given a perfect network *G*(*N, M*), where *N* and *M* are the nodes and link sets, calculate the node-level measure of interest *S*; (2) derive the rank *R*_*S*_ of *S*; (3) add normally distributed noise *ϵ* to the rank *R*_*S*_ such that the actual rank and the noisy rank *R*_*n*_ = rank(*R*_*S*_ + *ϵ*) are correlated by *ρ* ∈ (0, 1], where *ρ* is the Spearman rho rank correlation coefficient and *ϵ* ∼ 𝒩 (0, *σ*^2^); When *σ*^2^ = 0 and therefore *ρ* = 1, the actual rank and the noisy rank are precisely the same (perfect information). As *σ*^2^ → ∞ and *ρ* → 0, the noisy rank approaches a random ranking, which does not rely on network information. (4) Select the top-ranked nodes for vaccination based on the noisy rank *R*_*n*_.

We simulated the impact of increasing levels of network data error on the efficacy of network-based targeting of vaccination by investigating a range of *ρ* values from 1 (perfect network information) to 0 (no network information). Effectiveness was measured in terms of the relative difference in MNWIC when 20% of wards are vaccinated, at 75% within ward coverage, using the outcome of random vaccination as a baseline.

## 3 RESULTS

### 3.1 RVF virus transmission dynamics and the impact of cattle movements

At the end of 365 days in the absence of vaccination, our simulation model resulted in a mean cumulative incidence of infected wards of 79% in ER random networks, 75% in scale-free networks, 92% in small-world networks, and 74% in the data-driven network (Figure 4).

**Figure 4.**
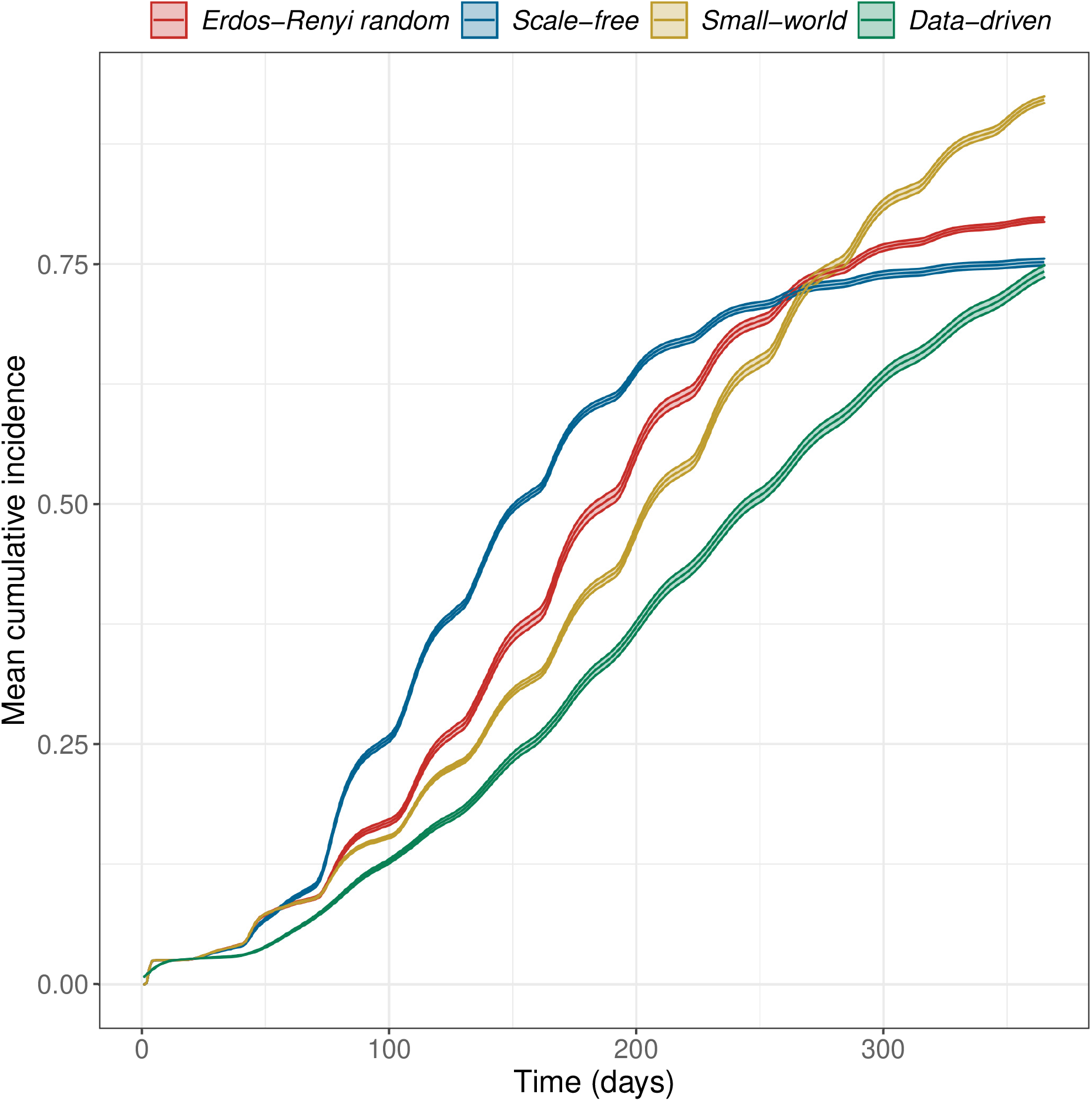
Simulated mean cumulative incidence by network type. The RVF meta-population model was simulated for one year, at a daily time step, with 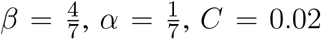 and 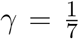, allowing disease spread within and between populations. Cattle movements between wards occur monthly, with rates determined by the weights of the data-driven network. In theoretical networks, 100 animals were moved between wards every month. This choice was based on the mean number of animals moved through the market network. The plots shown are the mean cumulative incidence of 300 simulations with a 95% confidence interval.

A larger outbreak (proportion of infected wards) was observed more often in small-world networks than in the other two theoretical network types, which can be explained through the size of the giant strongly connected components (GSCC) presented in Table 1. We observed the role of livestock movements on disease spread, occurring every month, through the sudden increase in cumulative incidence following the monthly movements (Figure 4).

### 3.2 Impact of intervention strategies

The effectiveness of targeted strategies varies with network type (Figure 5). Targeting vaccination at 10% of wards that rank highest on degree and betweenness reduced cumulative incidence by about 9% and 18% in ER random networks, 3% and 10% in scale-free networks, 15% and 22% in small-world networks, and 35% and 33% in the data-driven network. When 20% of wards are vaccinated, degree and betweenness strategies reduced cumulative incidence by over 50% in the data-driven network. At 20% degree and betweenness reduced cumulative incidence by about 33% and 47% in ER random networks, 11% and 24% in scale-free networks, 36% and 43% in small-world networks, and 59% and 58% in the data-driven network. Using a PageRank strategy, we need to vaccinate up to 40% of wards to reduce cumulative incidence by 50% in ER random and small-world networks; and just over 20% and 40% in data-driven and scale-free networks, respectively. Vaccination by PageRank showed better performance than degree vaccination in scale-free networks when the proportion of vaccinated wards is higher than 20%.

**Figure 5.**
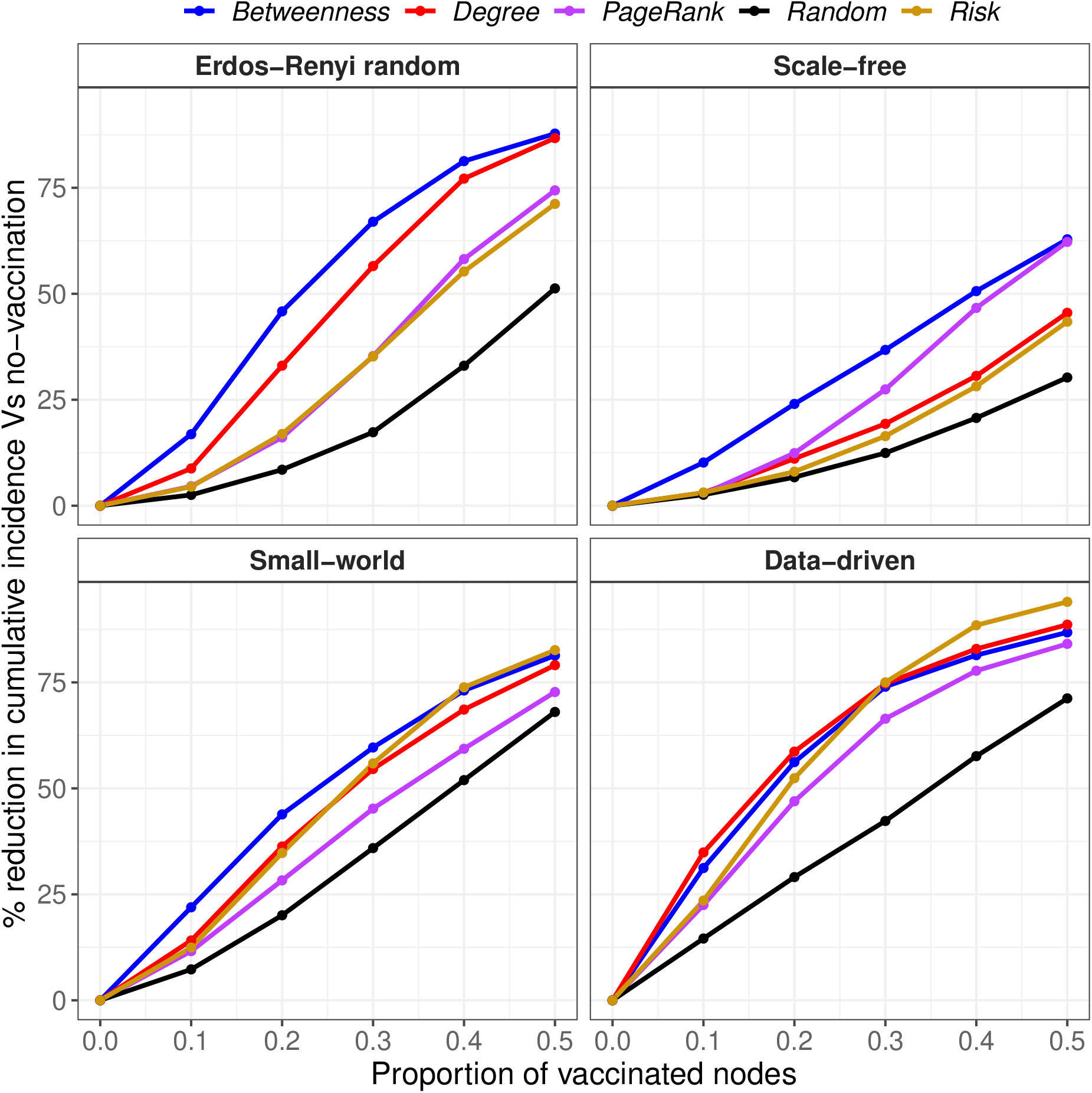
Percentage reduction in node cumulative incidence relative to no vaccination by the proportion of vaccinated nodes, at 75% within-node coverage, for different vaccination strategies.

In general, random vaccination offers the worst performance in all networks. The effectiveness of risk-based vaccination was not noticeably different from PageRank and degree in ER random and Scale-free networks, respectively. Our results show that vaccination by the risk of disease emergence is highly effective in the data-driven network. Only degree and betweenness were more effective when fewer than 30% of wards were vaccinated. Interestingly, it outperformed all network-based strategies when the proportion of vaccinated wards was 30% or higher.

### 3.3 Effect of adding noise on effectiveness

We examined the impact of incomplete network information on the effectiveness of vaccination strategies by adding noise to our assessment of risks. Figure 6 shows the relative difference in cumulative incidence when 20% of wards are vaccinated using random vaccination as a baseline. All targeted strategies outperformed random vaccination at all *ρ >* 0 across all networks. Even at a very high noise level (low values of *ρ*), targeted strategies reduced cumulative incidence further than random vaccination in all networks.

**Figure 6.**
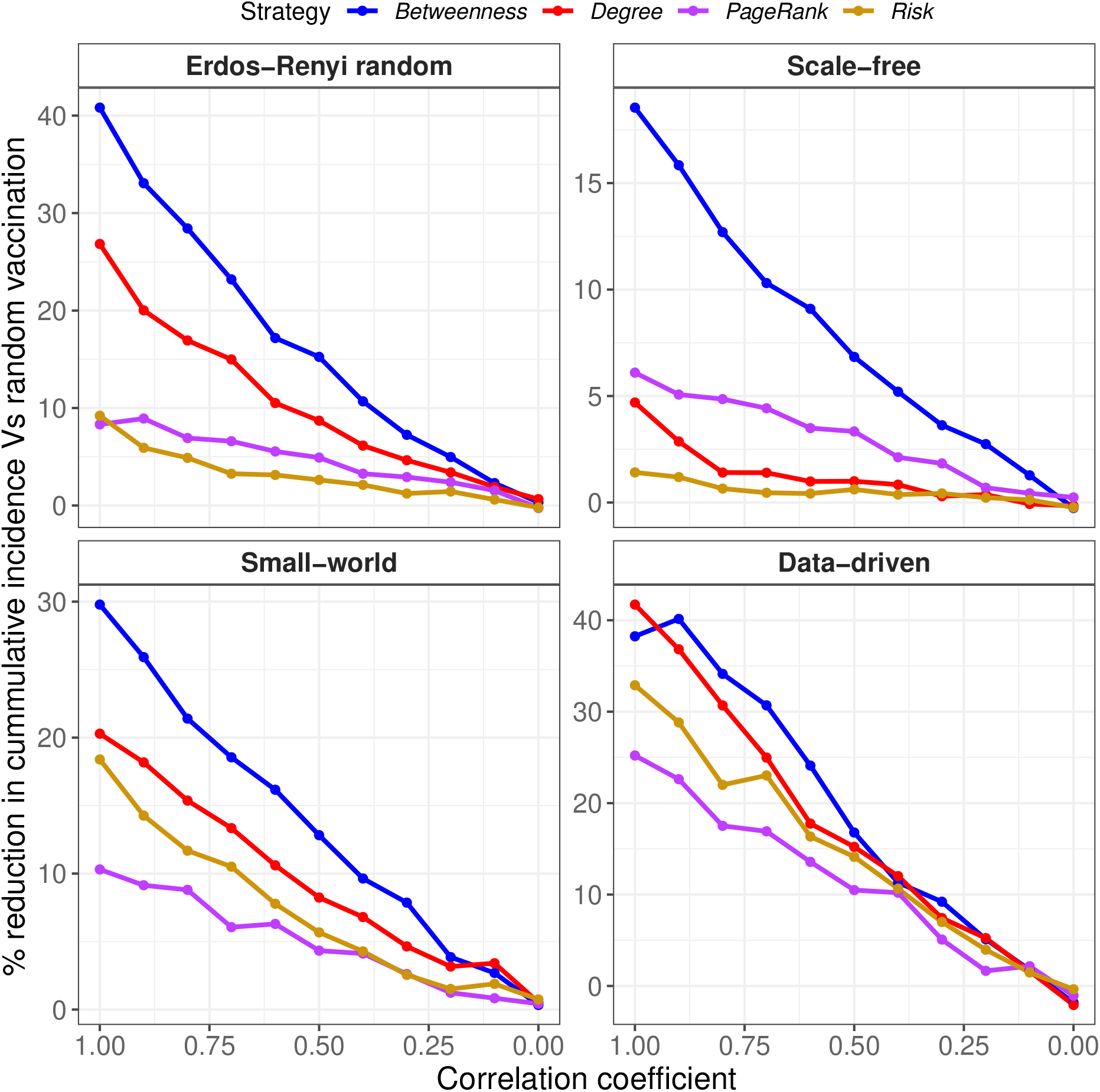
The effect of increasing noise on the effectiveness of vaccination strategies when 20% of nodes are vaccinated, at 75% within-node coverage. Noise was added to simulate conditions of imperfect network information. Perfect and imperfect ranks were compared using Spearman’s *ρ* rank correlation coefficient, with a correlation coefficient of 1 indicating perfect information.

The effectiveness of targeted strategies decreases almost linearly with increasing noise, and converges to random vaccination as *ρ* approaches 0. Similar trends were observed when 10%, 30%, 40%, and 50% of wards are vaccinated (see supplementary material).

When network information was perfect (*ρ* = 1), betweenness performed better than random vaccination by about 41% in ER random networks, 19% in scale-free networks, 30% in small-world networks, and 38% in the data-driven network (Fig. 6). Similarly, degree was better by about 27%, 5%, 20%, and 42% ER in random, scale-free, small-world, and data-driven networks, respectively. Betweenness and degree, both strategies lost their effectiveness by over 50% in all networks when our knowledge of network information dropped by half (*ρ* = 0.5). A similar observation was noted about risk strategy in the data-driven and small-world networks, where effectiveness drops by over 50% when our knowledge of disease emergence points is 50% correct. Even though vaccination by degree outperformed other strategies in the data-driven network at *ρ* = 1, their effectiveness was similar to risk-based vaccination when *ρ* was lower than 0.5.

## 4 DISCUSSION

Modelling techniques help describe disease dynamics and test the impact of interventions. In this study, we used a meta-population model to simulate the spread of RVF virus on livestock movement networks, examined the impact of network-based and risk-based vaccination strategies, and explored how imperfect information affects their effectiveness. This research addressed the common real-life problems in veterinary research — incomplete and patchy data about livestock movement patterns — arising from Tanzania and many other countries.

The major finding of this investigation is that the loss of effectiveness of targeted vaccination strategies with increasing data imperfection is approximately linear regardless of network structure. Essentially, any improvement in information reliability is beneficial.

Additional results showed no universally optimal targeting strategy across all network structures and proportions of vaccinated nodes. Nevertheless, vaccination by betweenness and degree are generally the most effective network-based strategies. This observation is robust to very different network structures and corroborates the results of other research studies (65, 66, 48, 29). Hébert-Dufresne et al. (48) used SIR and SIS models to compare the effectiveness of targeted immunisation strategies (degree, betweenness, community membership) on 17 networks, including World Wide Web, US power Grids and Co-authorship networks (MathSci and arXiv). Similarly, Eames et al. (66) studied the impact of preventive vaccination on a weighted contact network using a SIR model. Keeling et al. (29) compared pre-emptive vaccination with a combination of reactive vaccination and culling during an ongoing epidemic using a stochastic farm-based simulation model of foot-and-mouth disease in livestock. They found that vaccinating high-risk farms (based on the largest cattle farms) before an epidemic efficiently reduces the potential for a major outbreak. In our data-driven network, the effectiveness of betweenness and degree were similar when the proportion of vaccinated nodes was lower than 30%. Above 30%, degree vaccination dominates over betweenness. In this case, the choice of vaccination strategy depends on the resources available. Even though the data-driven network is fully connected with GSCC of 398, the outbreak size was smaller than that of the small-world with GSCC of 388 network. A possible explanation for this circumstance might be because up to 34% of the link weights in the data-driven network are significantly less than 1, representing lower onward transmission risk through those links.

Under perfect information, we noted a crossover point between the effectiveness of network-based (degree and betweenness) and risk-based vaccination in the data-driven network. For example, when a small proportion of wards are vaccinated (*<* 30%), both degree and betweenness vaccination work better than risk vaccination. However, when the proportion of vaccinated wards is higher, risk vaccination outperformed all network-based vaccination. This evidence suggests that if our knowledge of network information were perfect, there might be scenarios where vaccinating the points of RVF virus introduction into the system is preferred over vaccinating the points of onward transmission. In scenarios where we can only cover 20% of wards, our findings show there is a noise level at which the effectiveness of degree is no different from the effectiveness of risk vaccination. Again, this observation suggests that if our knowledge of network information is imperfect, depending on how well we know the highly central and high-risk wards, we might choose one strategy over the other when we can only vaccinate a few wards. Because this result is not robust to changing network structures, we are unable to extrapolate it to all networks, so it must be interpreted with caution. For instance, in theoretical networks, all targeting strategies are perfectly linear with increasing noise and there is no crossover.

There are some potential limitations of this study. First, the network used in this study is generated based on the outcome of Chaters et al. (24), and it is not an exact representation of the Tanzanian cattle movement system. Secondly, the number of nodes used to generate theoretical networks was kept very small to mimic the data-driven system, which might affect our ability to generalise results. Thirdly, we did not evaluate the impact of the bias and error associated with noise in the risk of RVF, as expressed through NDVI. Whilst NDVI can be accurately measured, its correlation with the emergence of RVF is not fully known and thus our strategy of targetting introduction into the highest NDVI wards may have relevant inaccuracies.

Lastly, our model did not consider mosquito-vector dynamics. This decision was due to insufficient data about the abundance and activities of specific mosquitoes (*Aedes* species) capable of transmitting the RVF virus. It is evident that mosquitoes contribute to keeping the RVF virus within the enzootic cycle by transmitting the virus from host to host, leading to the persistence of RVF. Our model can be extended to explicitly include mosquito population dynamics and multiple hosts where data are available. We recommend future studies on this topic, with additional complexities such as a detailed RVF virus transmission model, livestock movement system at a finer scale (e.g., village-level), and various intervention strategies (e.g., movement bans).

In conclusion, our study provides a framework to examine the hypothesis that the well-studied targeted strategies, which are highly effective under perfect information, might be ineffective under imperfect information. We hope this work will contribute to a new or existing framework for carefully designing and planning network-driven intervention strategies for infectious livestock disease control, particularly in sub-Saharan Africa, where missing livestock movement data severely limits opportunities for network-targeted interventions.

## AUTHOR CONTRIBUTIONS

TS, PJ, JE, SC, and RK conceptualised and formulated the model. TS wrote the simulation code, analysed the model, produced visualisations, interpreted results, and wrote the original draft of the manuscript. All authors reviewed the manuscript and approved the submitted version.

## FUNDING

This work was supported by the Biotechnology and Biological Sciences Research Council (BBSRC), the Foreign and Commonwealth Development Office (FCDO), the Economic & Social Research Council (ESRC), the Medical Research Council (MRC), the Natural Environment Research Council (NERC) and the Defence Science & Technology Laboratory, under the Zoonoses and Emerging Livestock Systems (ZELS) programme (BB/L018926/1). TS was supported by the University of Edinburgh & University of Glasgow Jointly Funded PhD Studentship in One Health. For the purpose of open access, the author has applied a Creative Commons Attribution (CC BY) licence to any Author Accepted Manuscript version arising from this submission.

## CONFLICT OF INTEREST STATEMENT

The authors declare that the research was conducted in the absence of any commercial or financial relationships that could be construed as a potential conflict of interest.

## ACKNOWLEDGMENTS

We are grateful to Divine Ekwem for his helpful discussion and input.

## DATA AVAILABILITY STATEMENT

The dataset used for this analysis records livestock movements in northern Tanzania recorded from 30 000 Livestock Movement Permits. This dataset belongs to the Ministry of Livestock and Fisheries of Tanzania, and we do not have permission to share it publicly. This also applies to the data on the number of cattle in each ward. Please contact the corresponding author for information on accessing this data.

## CODE AVAILABILITY STATEMENT

All code, including network generation, metapopulation model, and visualisation, are available in a public GitHub repository at https://github.com/tijanisulaimon/rvf_metapop_nTanzania.

## Notes

https://github.com/tijanisulaimon/rvf_metapop_nTanzania

### Competing Interest Statement

The authors have declared no competing interest.

