## Supplementary Figures for "Modelling the effectiveness of targeting Rift Valley fever virus vaccination using imperfect network information"

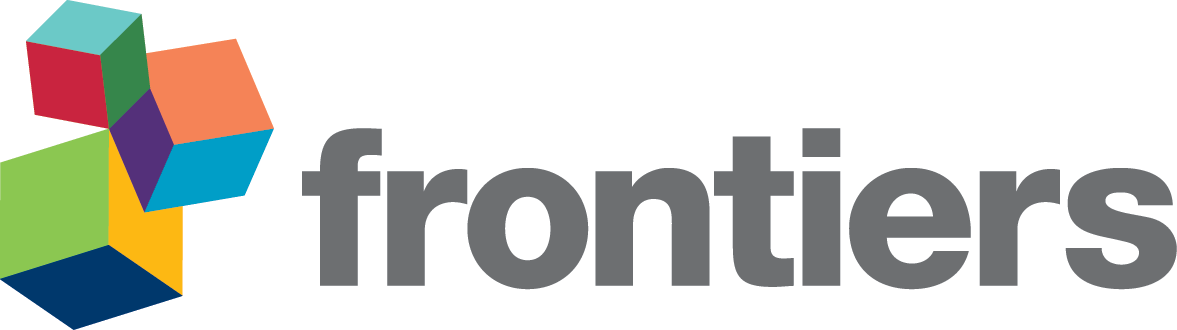

*Supplementary Material*

**1	SUPPLEMENTARY FIGURES**

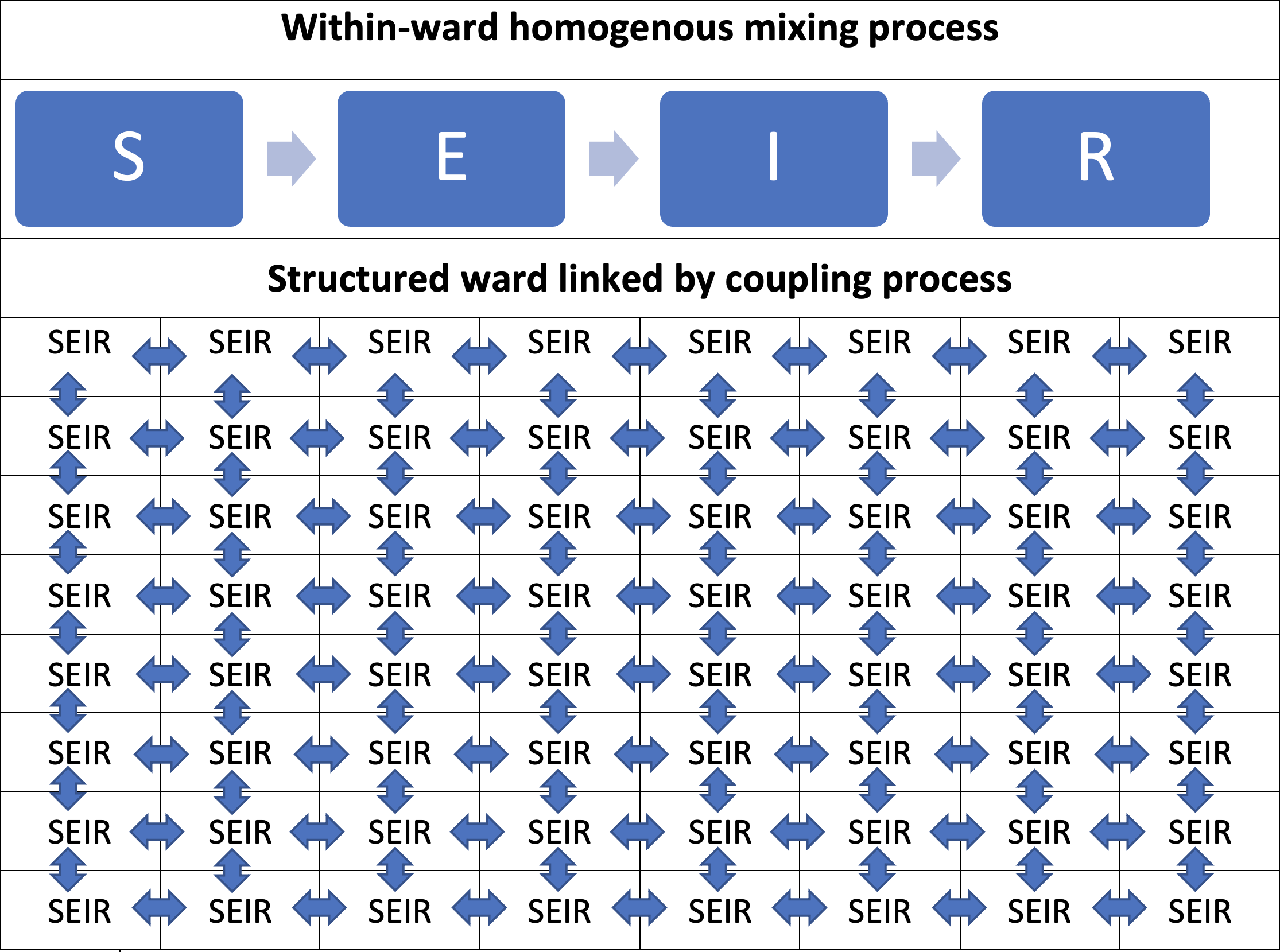

**Figure S1. Structured SEIR model describing transmission of Rift Valley fever virus within-wards in northern Tanzania**. Cattle movement data were available describing cattle movements between administrative wards in northern Tanzania and this is the fundamental unit of our data-driven network model. Each ward was divided into 8 8 gird cells to simulate local within-ward transmission driven by spatial transmission. A homogeneous mixing SEIR model was used to describe the transmission dynamics of RVF virus within each cell. Transmission between adjacent cells was allowed through spatial coupling.

*×*

The grey arrows represent the flow of cattle between four states disease states–susceptibles (S), exposed (E), infectious (I), and recovered (R).

***Supplementary Material***

*Betweenness	Degree	PageRank	Risk*

15	8

Erdos−Renyi random

Scale−free

6

% reduction in cummulative incidence Vs random vaccination

10

4

5

2

_0_	0

Small−world

Data−driven

25

15

20

10	15

10

5

5

0

1.00

0.75

0.50

0.25

0.00

0

1.00

0.75

0.50

0.25

0.00

### Correlation coefficient

**Figure S2.** The effect of increasing noise on the effectiveness of vaccination strategies when 10% of nodes are vaccinated, at 75% within-node coverage

***Supplementary Material***

Strategy	*Betweenness	Degree	PageRank	Risk*

60	30

Erdos−Renyi random

Scale−free

% reduction in cummulative incidence Vs random vaccination

40	20

20	10

50

Small−world

Data−driven

60

40	_50_

40

30

30

20

1.00

0.75

0.50

0.25

0.00

20

1.00

0.75

0.50

0.25

0.00

### Correlation coefficient

**Figure S3.** The effect of increasing noise on the effectiveness of vaccination strategies when 30% of nodes are vaccinated, at 75% within-node coverage

***Supplementary Material***

Strategy	*Betweenness	Degree	PageRank	Risk*

Erdos−Renyi random

| Scale−free |
| --- |

80

% reduction in cummulative incidence Vs random vaccination

40

60

30

40

20

80

Small−world

Data−driven

60	_70_

60

50

50

40

1.00

0.75

0.50

0.25

0.00

40

1.00

0.75

0.50

0.25

0.00

### Correlation coefficient

**Figure S4.** The effect of increasing noise on the effectiveness of vaccination strategies when 40% of nodes are vaccinated, at 75% within-node coverage

***Supplementary Material***

Strategy	*Betweenness	Degree	PageRank	Risk*

60

Erdos−Renyi random

Scale−free

80

% reduction in cummulative incidence Vs random vaccination

50

70

40

60

_50_	30

90

Small−world

Data−driven

75

80

70

65	70

60	_60_

1.00

0.75

0.50

0.25

0.00

1.00

0.75

0.50

0.25

0.00

### Correlation coefficient

**Figure S5.** The effect of increasing noise on the effectiveness of vaccination strategies when 50% of nodes are vaccinated, at 75% within-node coverage
